# Mutations in Complex I of the mitochondrial electron-transport chain sensitize the fruit fly (*Drosophila melano*gaster) to ether and non-ether volatile anesthetics

**DOI:** 10.1101/2022.11.22.517557

**Authors:** Luke Borchardt, Amanda R. Scharenbrock, Zachariah P. G. Olufs, David A. Wassarman, Misha Perouansky

## Abstract

The mitochondrial electron transport chain (mETC) contains molecular targets of volatile general anesthetics (VGAs), which places carriers of mutations at risk for anesthetic complications. The *ND-23^60114^* and *mt:ND2^del1^* lines of fruit flies (*Drosophila melanogaster*) that carry mutations in core subunits of Complex I of the mETC, replicate numerous characteristics of Leigh syndrome (LS) caused by orthologous mutations in mammals and serve as models of LS. *ND-23^60114^* flies are behaviorally hypersensitive to volatile anesthetic ethers and develop an age- and oxygen-dependent anesthetic-induced neurotoxicity (AiN) phenotype after exposure to isoflurane but not to the related anesthetic sevoflurane. The goal of this paper was to investigate whether the alkane volatile anesthetic halothane and other mutations in Complex I and in Complexes II-V of the mETC cause AiN. We found that (i) *ND-23^60114^* and *mt:ND2^del1^* were susceptible to toxicity from halothane; (ii) in wildtype flies halothane was toxic under anoxic conditions; (iii) alleles of accessory subunits of Complex I predisposed to AiN; and (iv) mutations in Complexes II-V did not result in an AiN phenotype. We conclude AiN is neither limited to ether anesthetics nor exclusive to mutations in core subunits of Complex I.

**Previous presentations:** parts of the data were presented in abstract form at the 63^rd^ Annual Drosophila Research Conference in San Diego, March 2022.

## 1. Introduction

Anesthetic-induced neurotoxicity (AiN) is a potential concern anytime volatile general anesthetics (VGAs) are administered, but certain conditions increase the risk of AiN. Risk factors include the extremes of age, the fragile and/or diseased brain and genetic factors.[1] Among the most relevant risk factors for ‘collateral’ effects of VGAs are interactions with mitochondrial function. The mitochondrial electron transport chain (mETC) contains molecular targets of VGAs and mETC function is dose-dependently suppressed by VGAs at clinically relevant concentrations in ‘wildtype’ mitochondria.[2] Mutations in Complex I of the mETC increase the sensitivity of oxidative phosphorylation to depression by VGAs resulting in anesthetic hypersensitivity [3] and possibly of increased risk for AiN.[4, 5] Over 80 mutations mostly affecting Complex I have been identified as causing Leigh Syndrome (LS), a rare incurable neurodegenerative disease resulting in severe disability and early death and associated with increased perioperative morbidity and sensitivity to VGAs.[6]

Although LS is a risk factor for perioperative complications, LS patients are frequently exposed to anesthesia for diagnostic and surgical procedures.[4] Animal models of LS reproduce its key pathological characteristics including hypersensitivity to behavioral effects of VGAs that can be used for a better understanding of risk factors for AiN.[7–9] The *ND-23^60114^* and *mt:ND2^del1^* lines of fruit flies (*Drosophila melanogaster*) are models of LS caused by hypomorphic mutations in the nuclearly encoded ND-23 (mammalian Ndufs8) and the mitochondrially encoded mt:ND2 (mammalian ND2) subunits that are conserved proteins of the core of Complex I.

Young adult *ND-23^60114^* flies present with a phenotype of increased behavioral sensitivity to the VGAs isoflurane (ISO) and sevoflurane (SEVO) replicating mammalian data.[10] At 10-13 days old, *ND-23^60114^* flies develop a lethal phenotype within 24 h after exposure to ISO but not to SEVO.[11] AiN is the likely cause because this phenotype is rescued by neuron-specific overexpression of wildtype ND-23.[11] Interestingly, heterozygous *ND-23^60114^* flies are asymptomatic and have a normal lifespan indicating haplosufficiency of ND-23.[8] However, at 30-35 days of age, *ND-23^60114^*/+ flies develop an AiN-like lethal phenotype after exposure to ISO in 75% O_2_ (hyperoxic ISO), indicating that effects of aging may result in haploinsufficiency of the core Complex I subunit under environmental stress conditions.[11]

The present study was designed to test the hypotheses that (i) the VGA halothane (HAL, an alkane as opposed to the ethers ISO and SEVO) causes AiN; and (ii) previously not examined mutations in nuclearly encoded genes of Complexes I-V are associated with AiN.

Our results indicate that HAL is toxic to the LS-model mutants in subunits ND-23 and mt:ND2. We found that mutation of the accessory subunit ND-SGDH (mammalian NDUFB5) is associated with a lethal phenotype after exposure to hyperoxic ISO. We found that mutations in Complexes II-V did not cause mortality following exposure to hyperoxic ISO. We conclude that mutations in Complex I impart the highest risk for AiN and the risk extends to mutations in accessory (*i.e*., non-core) subunits. We also discovered that HAL sensitizes wildtype Canton S flies to anoxia and that AiN is only loosely linked to increased intestinal permeability (IP).

## 2. Results

### 2.1. *Mutants of Complex I subunits* ND-23 *and* mt:ND2 are sensitive to halothane toxicity

We exposed mixed sex, 11 to 13 day-old Canton S (wildtype), *ND-23^60114^* and *mt:ND2^del1^* flies to 2 h of 1.5% HAL in room air). HAL did not affect mortality in 21% and 75% O_2_ in Canton S flies. In contrast, exposure to HAL increased mortality from a natural attrition rate of 6.8±1.64% and 25.9±3.07% in 21% O_2_ to 21.7±4.97% and 56.6±3.11% in *ND-23^60114^* and *mt:ND2^del1^* flies, respectively (p<0.0001 for both genotypes, unpaired t-test, Fig 1). Hyperoxia (75% O_2_) increased mortality from HAL in *ND-23^60114^* flies but but the difference to mortality in normoxia was not significant (p<0.16, unpaired t-test). However, hyperoxia increased HAL-mortality in the *mt:ND2^del1^* line from 56.6±3.11% to 96.4±1.11% (p<0.0001, unpaired t-test). These results indicate that HAL is toxic to mitochondrial mutants. Because previous reports showed that ISO-induced AiN in *ND-23^60114^* flies was suppressed by hypoxia, we tested HAL-toxicity in 5% O_2_. However, hypoxia did not significantly suppress mortality following HAL exposure in either *ND-23^60114^* or the *mt:ND2^del1^* flies. Finally, mortality resulting from 2 h of anoxia in *ND-23^60114^* and *mt:ND2^del1^* flies was not altered by HAL, but it increased mortality in Canton S flies from 8.1±1.52% to 26.6±6.37% (p=0.018, unpaired t-test). We conclude that HAL, while chemically different from ISO, is associated with AiN in mutants of core Complex I subunits. Furthermore, HAL also is toxic under anoxic conditions in wildtype Canton S flies.

**Figure 1.**
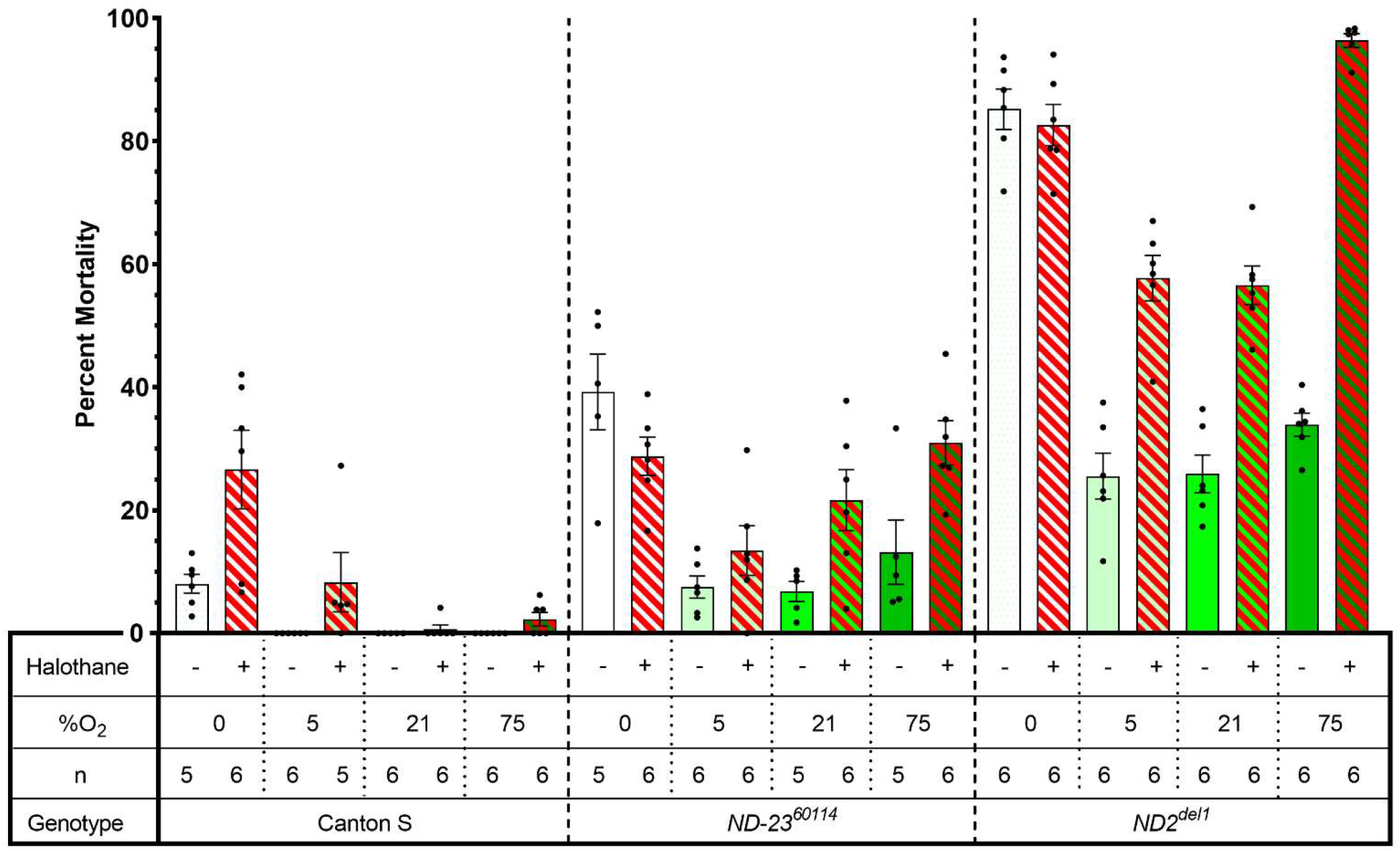
Halothane causes AiN in mitochondrial mutants. 11 to 13 day-old flies of three genotypes were exposed to two hours of 1.5% HAL administered by a carrier gas containing between zero and 75% O_2_ and corresponding concentrations of N_2_. The number of dead flies was counted 24 h after exposure. Data shown as mean ±SEM, n indicates number of biological replicates of 25-30 mixed sex flies each shown as dots. Note: high rates of natural attrition in mutants.

### 2.2. Complex I: amutant allele of ND-SGDH results in AiN

Complex I (NADH-ubiquinone oxidoreductase) is with 1MDa and 44 or 45 subunits the largest complex of the mETC consisting of a peripheral and a membrane arm. Fourteen core subunits contain the catalytic machinery. These are sufficient for oxidoreductase function and are conserved from bacteria to mammals. The remaining subunits (31-32 in mammals, 28 in *Drosophila*) are accessory and phylum-specific. Complex I is L-shaped with a membrane arm and a matrix arm. [12] About 30% of mitochondrial diseases affecting energy metabolism are caused by mutations in the nuclearly of mitochondrially encoded subunits of Complex I. [13] Some tested mutations sensitize animals to the behavioral effects of VGAs and may represent perioperative risk factors.

*ND-23^60114^* flies showed a strong AiN phenotype at 2 weeks (8-13) days old but not earlier (Fig 2A, A’), while Canton S flies showed no AiN throughout the tested lifespan (Fig 2A).

**Figure 2.**
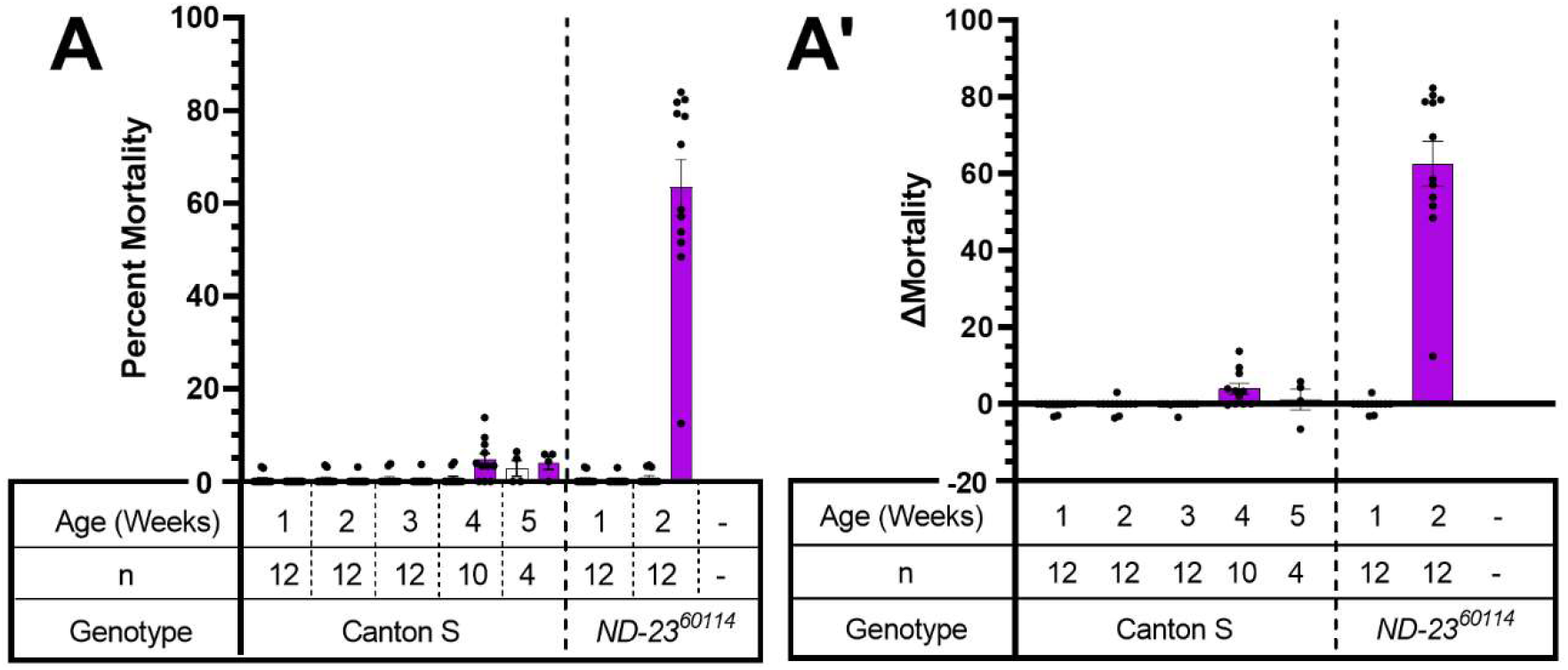
AiN in *ND-23^60114^* flies is age-dependent. Flies were exposed to two hours to 2% ISO in 75% O_2_ at weekly intervals beginning at one week of age and returned to standard culture vials. 24 h after exposure, dead flies were counted and survivors transferred to fresh vials. No mortality was observed in Canton S flies while *ND-23^60114^* experienced an abrupt increase in mortality at two weeks of age (Panel A). Panel A’: same data as A with control mortality subtracted (□ Mortality). Each dot represents a biological replicate of 25-30 mixed sex flies. Results are presented as mean± SEM

To investigate the importance of other Complex I subunits for VGA toxicity, we screened the mutants listed in Table I. All lines are homozygous viable and were tested under identical conditions at multiple time points during their lifespan (i.e., survivors after each exposure to hyperoxic ISO were maintained under standard culture conditions and exposed again to hyperoxic ISO at the subsequent time point until their number fell below the preset threshold of four vials considered the minimum for meaningful analysis (Fig. 3).

**Table I.**
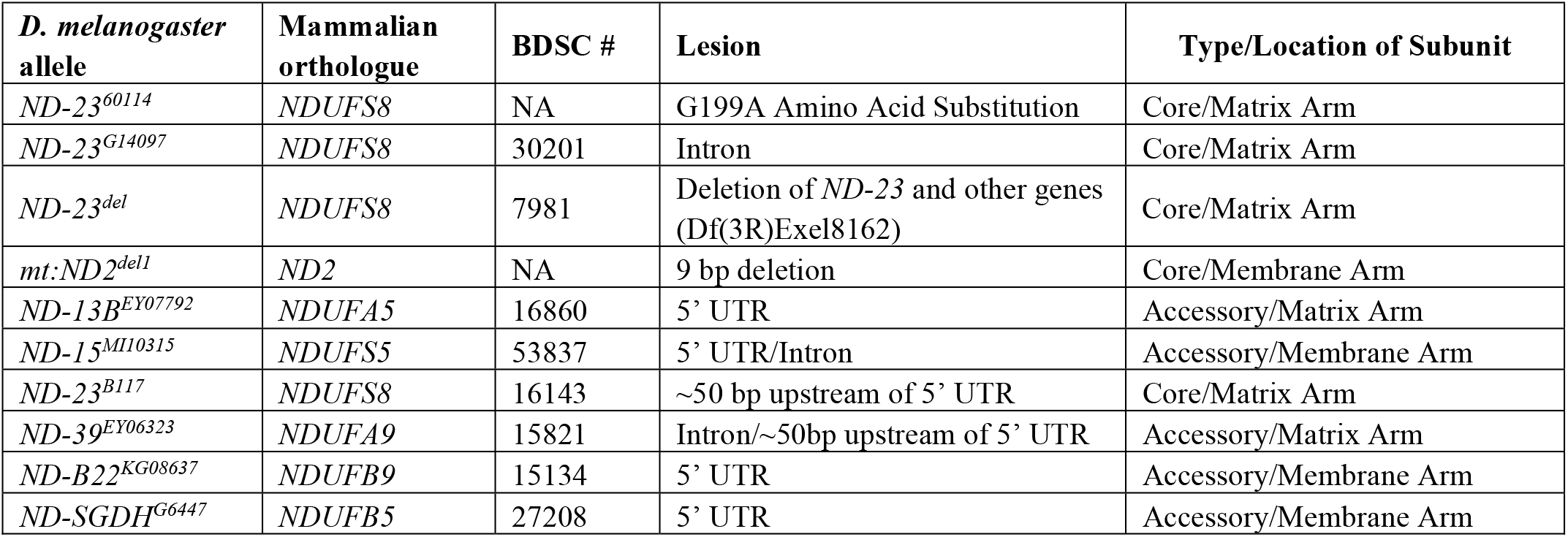
Complex I alleles screened for hyperoxic ISO toxicity. BDSC: Bloomington Drosophila Stock Center.

**Figure 3.**
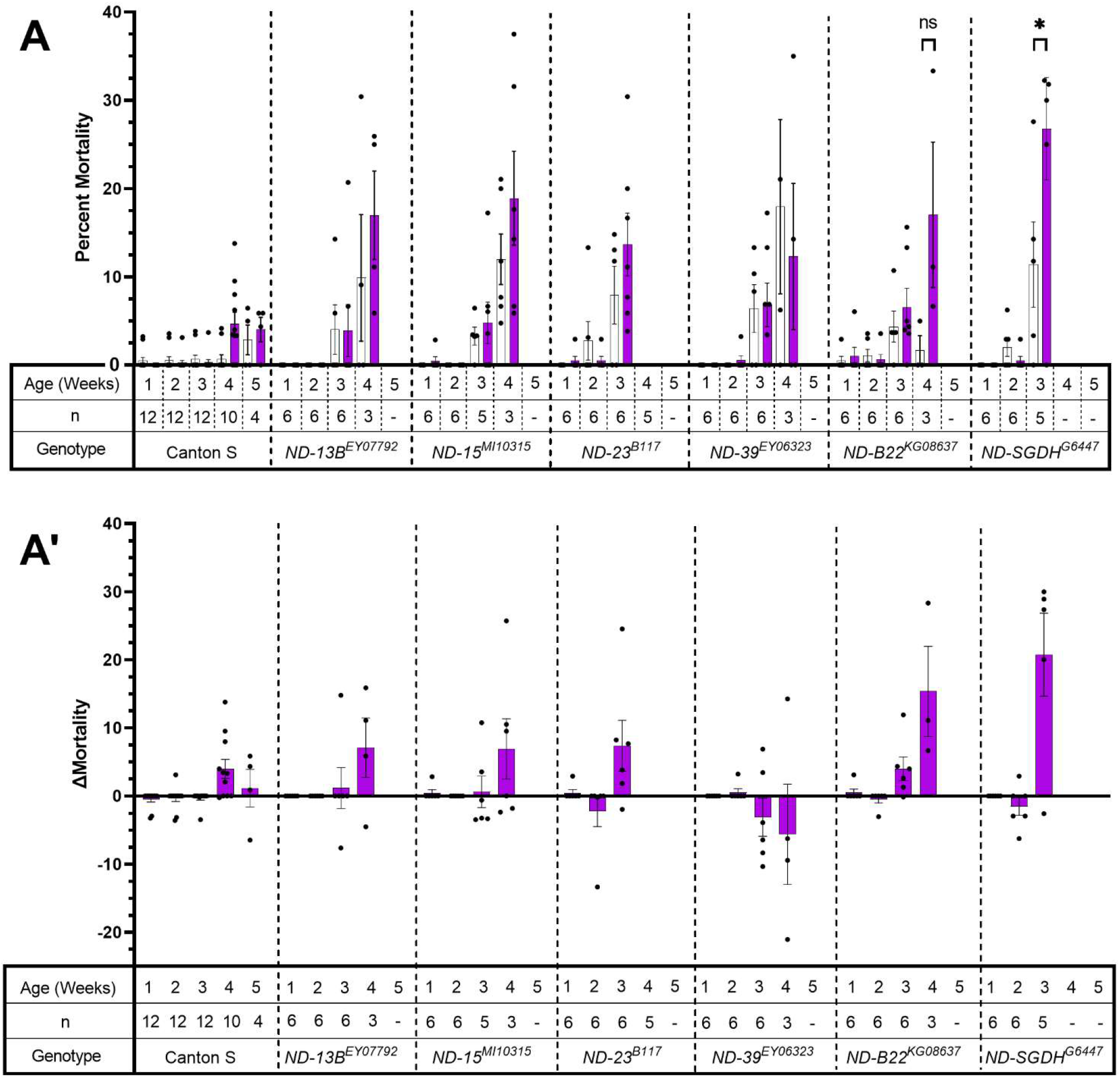
Complex I subunits screen identifies an *ND-SGDH* mutant as AiN susceptible. A. Flies were exposed for two hours to 2% ISO in 75% O_2_ at weekly intervals beginning at one week of age. After exposure, they were returned to standard culture vials. 24 h after exposure, dead flies were counted and survivors were transferred to fresh vials. Mutations in Complex I subunits carried variable age-dependent mortality due to natural attrition (white bars). The increase in mortality after hyperoxic ISO reached significance only for *ND-SGDH^G6447^* at three weeks old. A’. Data from A, control mortality subtracted (□ Mortality). (Week 1 = 1-6 days old; Week 2 = 8-13 days old; Week 3 = 15=20 days old; Week 4 = 22-27 days old; Week 5 = 29-34 days old). Each dot represents a biological replicate of 25-30 mixed sex flies. Results are presented as mean± SEM.

Mortality of *ND-SGDH^G5447^* mutants was increased by hyperoxic ISO from 11.4±4.82% to 26.8±5.80% (p=0.027, one-way paired t-test). For *ND-B22^KG08637^* mortality at 4 weeks also appears increased (but lacked statistical significance p= 0.073 one-way paired t-test) which might be due to the low number of biological replicates available at this age. To confirm the positive finding of the screen (Fig 3), we re-tested *ND-SGDH^G5447^*) at a single time point 15-20 days of age (Fig 4). Mortality 24 h after exposure to hyperoxic ISO was increased by 12.8±3.16% (from 38.69±3.16% to 51.49±3.23%; p=0.0254, unpaired t-test). We conclude that mutations in accessory subunits of Complex I can increase the risk of AiN. At the same time, the age at which AiN manifests may vary between different alleles of the same gene. For example, the *ND-23^60114^* allele shows early severe AiN while *ND-23B^117^* had increased (but not statistically significant) mortality at 3 weeks.

**Figure 4.**
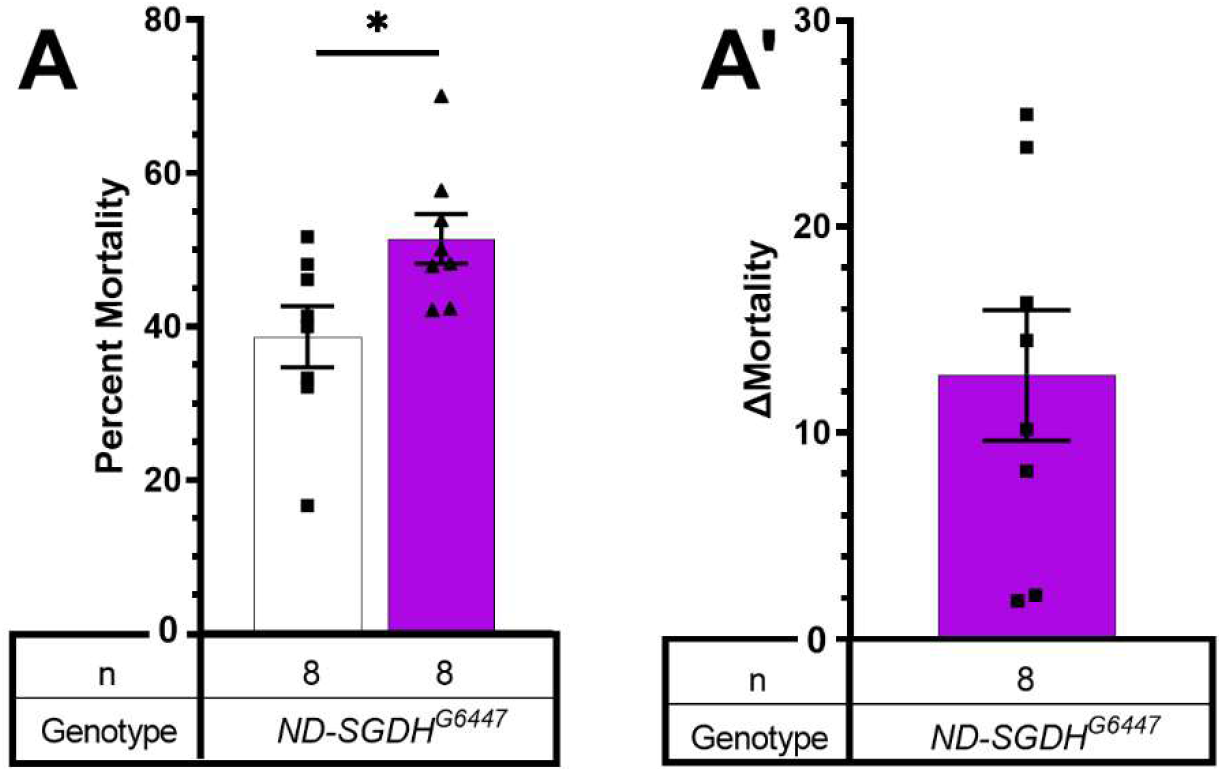
Hyperoxic ISO increases 24 h mortality in *ND-SGDH^G6447^* mutants. A. Flies were exposed to 2% ISO in 75% O2 for two hours at 15-20 days old, returned to culture vials and scored 24 h after exposure. Mortality increased in mutants but not in wildtype flies (p=0.0254, unpaired t-test). A’. Data as in A, control mortality subtracted (□ Mortality). Each dot represents a biological replicate of 25-30 mixed sex flies. Results presented as mean±SEM.

### 2.3. The tested Complex II-V mutants did not show AiN

To investigate a potential role of mETC Complexes II-V in VGA toxicity, we screened mutants carrying homozygous viable alleles in Complexes II-V listed in Table II (Fig. 5). Additionally, we tested one Complex II (Succinate dehydrogenase □-subunit, SdhB) allele (SdhB^ey12061^/CyO) as a heterozygote because it is homozygous lethal. All lines were tested under identical conditions repeatedly (i.e., survivors after each exposure were exposed again at the subsequent time point until their number fell below a preset threshold considered as the minimum for meaningful analysis (see Materials and Methods)). None of the mutants had significantly increased mortality from exposure to hyperoxic ISO (Fig 5 A, A’). We conclude that homozygous viable alleles of Complexes II-V do not incur AiN even under hyperoxic conditions and at an advanced age.

**Table II.**
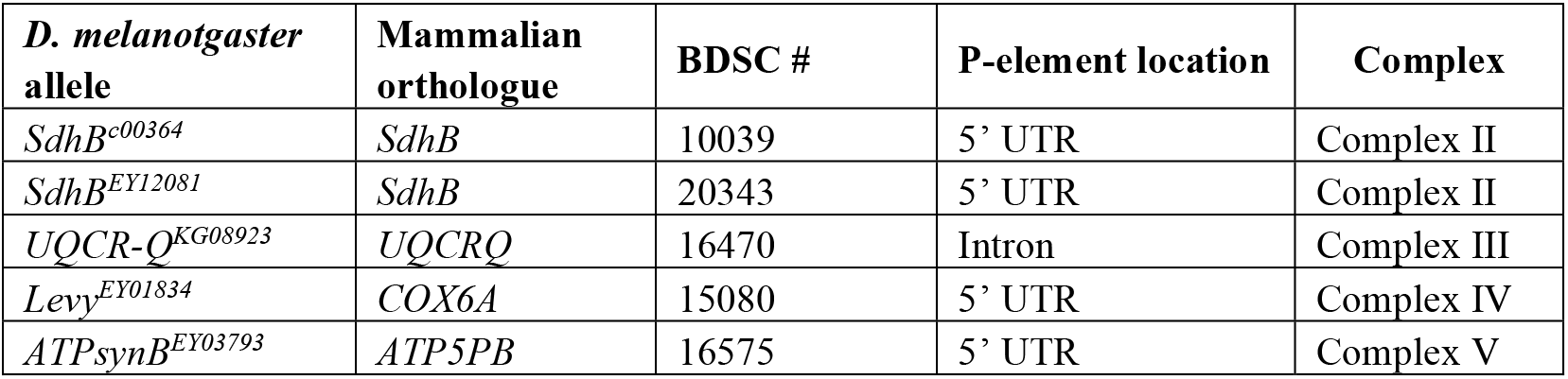
Alleles of Complexes II-V screened for hyperoxic ISO toxicity. BDSC: Bloomington Drosophila Stock Center.

**Figure 5.**
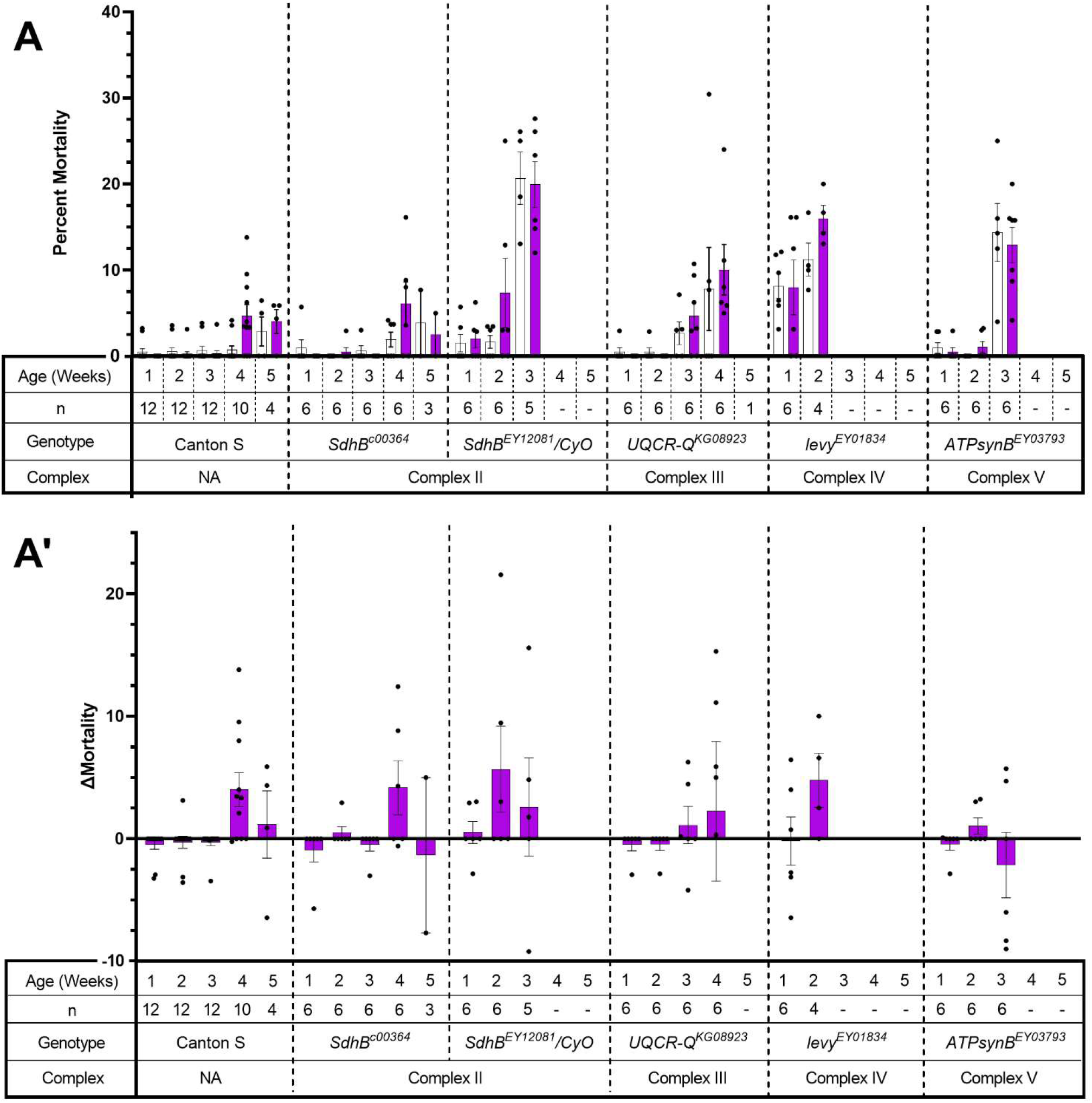
Hyperoxic ISO does not increase 24 h mortality in mutants of Complexes IIV. A. Repeated exposure at weekly intervals for two hours to 2% ISO in 75% O_2_ at 1-5 weeks of age. 24 h after exposure, dead flies were counted and survivors were transferred to fresh vials. Results presented as mean ± SEM. A’. Data as in A with control mortality subtracted (□ Mortality). Each dot represents a biological replicate of 25-30 mixed sex flies. (Week 1 = 1-6 days old; Week 2 = 8-13 days old; Week 3 = 15=20 days old; Week 4 = 22-27 days old; Week 5 = 29-34 days old)

### 2.4. AiN is not associated with increased intestinal permeability

Traumatic brain injury (TBI) causes increased intestinal permeability (IP) in mammals and flies.[14, 15] Increased IP can be easily assessed in flies by the ‘smurfing’ assay. [16] In uninjured, aging flies, increased IP is a harbinger of impending death. [17] In the fly TBI model, smurfing is highly predictive of mortality within 24 h after injury.[15] To test the degree to which mortality from AiN is associated with changes in IP, we fed blue-colored food to mixed sex wildtype Canton S and to *ND-23^60114^* flies 10-13 days of age prior to exposing them to ISO. Canton S flies neither smurfed nor died during the observation period (Fig. 6). Background mortality in *ND-23^60114^* control group was 6%, smurfing occurred 0.3%, which is similar to the 0.7% spontaneous smurfing rate reported previously in <u>*w^1118^*</u> flies[15]). After exposure to ISO, 6% of flies smurfed (p=0.0018) and 40.4% died (i.e., exposure to ISO increased IP but the majority of flies died from AiN without an increase in IP) (Fig. 6). These data suggest that signaling to the intestine originating in the brain that leads to increased IP is not an obligatory contributor to AiN.

**Figure 6.**
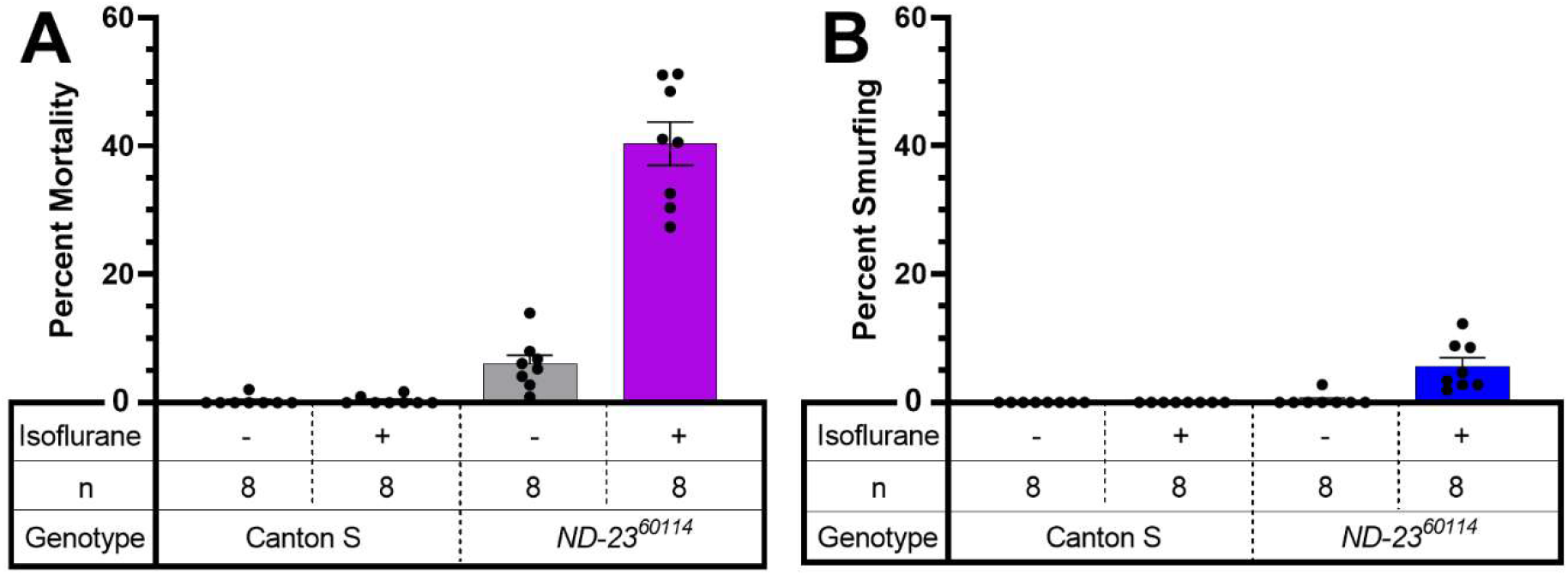
Mortality from AiN is not a result of increased intestinal permeability (IP). A. Normoxic ISO increases mortality 24 h after exposure in *ND-23^60114^* but not in wildtype flies. Note: unexposed *ND-23^60114^* incur a background mortality over the obserbvation period. B. No Canton S flies showed increased IP after exposure to ISO but a small fraction of *ND-23^60114^* flies (close to the background mortality without ISO exposure) smurfed. 10-13 day-old flies were fed a nonabsorbable blue dye for 48 h prior to exposure. Dead and smurfed flies were counted 24 h after exposure. Each dot represents a biological replicate of 25-30 mixed sex flies. Results presented as mean± SEM

## 3. Discussion

The principal findings presented in this manuscript are: (i) the alkane HAL causes AiN in flies harboring mutations of core Complex I subunits ; (ii) HAL sensitizes wildtype (Canton S) but not mutant (*ND-23^60114^* and *mt:ND2^del1^*) flies to anoxic injury; (iii) mutations in accessory Complex I subunits are associated with hyperoxic ISO-AiN; (iv) mutations in Complexes II-V are not associated with hyperoxic ISO-AiN; (iii) death from AiN is not associated with increased intestinal permeabilty (IP).

### 3.1. HAL causes AiN in mitochondrial mutants

Most mammalian data on AiN and our previous findings in the fly were obtained using the ether ISO. The current experiments aimed to answer the question whether AiN in mitochondrial mutants is ISO-specific. Because work in worms suggests that the behavioral sensitivity of the alkane VGA HAL is affected by mitochondrial mutations similarly to that of ISO,[2] we used HAL in two mutant fly lines carrying either the mitochondrially-encoded mt: *ND2^del1^* allele or the nuclearly-encoded *ND-23^60114^* allele. HAL differs from ISO in a number of ways: in contrast to the pungent ether ISO (2-Chloro-2-(difluoromethoxy)-1,1,1-trifluoro-ethane), HAL (2-Bromo-2-chloro-1,1,1-trifluoroethane) is a non-pungent alkane with a lower degree of fluorination and higher lipid solubility. In mammals, HAL undergoes a higher degree of metabolic transformation than ISO.[18, 19] Our findings of increased mortality after exposure to HAL in both *ND-23^60114^* and *mt:ND2^del1^* flies indicate that HAL is toxic to flies carrying mutations in the core of Complex I.

Previously, we showed that ISO toxicity in the *ND-23^60114^* model is caused by injury to the brain and is not specific to the *ND-23^60114^* allele.[11] We hypothesize that mortality from HAL is also caused by AiN because of the phenotypic similarity: the dose of HAL was equipotent to that of ISO, death occurred within 24 h after initial recovery from the anesthetic and mortality was modulated by the O_2_ concentration. Furthermore, neither HAL nor ISO are toxic to Canton S flies under normoxic and hyperoxic conditions. Interestingly, HAL increased mortality in Canton S under hypoxic/anoxic conditions which is reminiscent of hepatic halothane toxicity in mammals.[20, 21] Unexpectedly, HAL did not increase mortality under hypoxic or anoxic conditions in mutant flies, indicating that the significance of HAL’s interaction for toxicity may depend on the integrity of mETC Complex I.

### 3.2. Mutations in accessory subunits sensitize to AiN

In mammals, a mutation in the accessory *NDUFS4* subunit underlies a mouse model of LS and increases behavioral sensitivity to VGAs.[22] Therefore, we hypothesized that accessory subunits may carry an AiN phenotype. Viable homozygous mutants of *ND-18*, the fly orthologue of *NDUFS4*, are not available hence we selected subunits that were not previously tested for a behavioral anesthetic phenotype.[23] While we did not screen all available alleles of the 46 subunits comprising Complex I, we provide a ‘proof of concept’ by identifying one accessory subunit of Complex I that causes AiN. The *ND-SGDH^G6447^* allele significantly increased mortality at 15-20 days of age (Fig.3, 4). On the other hand, the *ND-23B^117^* allele showed a weaker (not reaching statistical significance) phenotype at a more advanced age than *ND-23^60114^* (Fig. 2 and 3). *ND-23B^117^* is caused by a P-element insertion ~ 50 bp upstream of the 5’-UTR probably resulting in a weaker ND-23 allele than the non-synonymous G199A polymorphism resulting in an amino acid substitution at position 19 that underlies the hypomorphic *ND-23^60114^* phenotype. These experiments suggest that the pharmacodynamic impact of a mutation is determined not only by the subunit and the functional significance of the specific allele: age and environmental conditions can modify the severity and time of onset of the disease phenotype as observed in Leber’s hereditary optic neuropathy.[24]

### 3.3. Mutations in Complexes II-V do not result in AiN

Previous work in *C. elegans* indicated that behavioral hypersensitivity to VGAs was not affected by mutations in the mETC unless they affected Complex I and that sensitivity to VGAs correlated with Complex I-dependent oxidative phosphorylation.[23] Our findings using AiN as an endpoint in mutations of subunits not tested by Falk et al. are generally in agreement with their findings: disruption of Complex I function is the determinant for adverse interactions with VGAs. A major limitation of these results is that we tested only single alleles. However, the tested mutations were functionally significant as illustrated by the high natural attrition and limited lifespan of all tested mutants.

In summary, mutations in various Complex I subunits carry the risk of AiN. Our data also indicates that age and environmental conditions influence the phenotypic presentation. Within the stated limitations (see 3.5), mutations in Complexes II-V do not appear to be associated with AiN.

### 3.4. AiN differs from TBI in its systemic impact

It was previously shown that AiN by ISO in *ND-23^60114^* flies was caused primarily by injury to the brain.[11] We wondered whether the nature of the toxic insult had commonalities with the systemic pathophysiology associated with TBI specifically with the effect of TBI on IP. At a 24 h mortality level of ±25% after experimental TBI in the <u>*w^1118^*</u> line (a standard laboratory fly strain)[15] that is comparable to mortality from AiN in *ND-23^60114^* flies, smurfing and death correlated tightly. We did not see such a correlation between AiN, IP and death (Fig. 6). The increase in IP after exposure to ISO may be due to local effects of ISO on intestinal epithelial tight junctions, analogous to its effect on the blood-brain barrier [25] or to a non-specific toxic effect but it is unlikely to be a direct consequence of AiN.

### 3.5. Limitations

We used behaviorally equipotent concentrations of HAL and ISO which may not be toxicologically equipotent. We also avoided quantitative comparisons of AiN-susceptibility across genotypes because of important confounding parameters: the highly different rates of natural attrition between genotypes, variability in mortality and, specifically for experiments involving HAL (Fig. 1) the different effects of O_2_ concentration. We also did not adjust experimental timing to lifespans: as a result, experiments were conducted at a more advanced biological age in mutants than in Canton S which may affect the penetrance of AiN.

## 4. Materials and Methods

The manuscript adheres to the applicable ARRIVE (Animal Research: Reporting of In Vivo Experiments) reporting guidelines (preclinical animal research). Approval from the Institutional Animal Care and Use Committee has been waived.

### 4.1. Fly Lines and Culturing

All flies were cultured at 25°C on cornmeal molasses food. See Tables 1 and 2 for information on fly lines used. Fly lines were obtained from the Bloomington Drosophila Stock Center, except for Canton S, which is our laboratory strain, and *ND-23^60114^*, which was provided by Barry Ganetzky (UW-Madison) [8] and mt: *ND2* which was a kind gift from the L. Pallanck (University of Washington, Seattle) lab [26] Flies were tested as homozygotes except where noted in figure legends.

### 4.2. Anesthesia and O_2_ Exposure

Flies were raised at 25°C for to two or five days post ecclosion then transferred to vials of 35 mixed-sex flies using CO_2_ and placed at 29°C. Vials were placed on their sides to prevent flies becoming stuck in the food. Flies were transferred to fresh food vials three times per week. On the day of exposure, flies were transferred to 50 mL conical tubes without use of CO_2_ and placed onto the serial anesthesia array (SAA) [10]. A mixture of N_2_ or O_2_, and air (21% O_2_, 79% N_2_) (Airgas, USA) were used to generate hypoxic (5% O_2_) or hyperoxic (75% O_2_) conditions using a clinical anesthesia machine (Aestiva/5, Datex-Ohemda) and confirmed by an inline internal O_2_ sensor. N_2_ alone was used to generate anoxic (0% O_2_) conditions. 1.5% HAL or 2% ISO were delivered by commercial agent-specific vaporizers (Ohmeda Fluotec 4 for HAL and Ohmeda Isotec 5 for ISO). After exposure to the desired conditions for two hours, the SAA was flushed with air for 5 minutes. Flies were transferred to fresh food culture vials, and incubated at 29°C for 24 hours. Percent mortality was determined by counting the number of dead flies and dividing by the total number of flies per vial. Mortality is a binary readout and was assessed by an unblinded observer. For multiple-exposure experiments, surviving flies were placed on fresh food vials at 29°C after mortality was scored until the next exposure. Multi-exposed flies were tested at weekly intervals. When fewer than 15 flies survived in a given vial, the vial was discarded. Experiments on a specific fly line were terminated when fewer than four vials remained. Unexposed flies were handled as above but were not placed on the SAA. Each experiment was conducted on at least three different days.

### 4.3. Smurfing

Flies were fed a nonabsorbable blue dye (FD&C blue dye no. 1©, Spectrum Chemical MFG Corp.) at a concentration of 2.5% in standard food. Flies were kept overnight at 29°C, visually inspected for the presence of blue color and excluded from the experiment if no color was seen. They were then exposed to 2% ISO for two hours. Following the exposure, flies were placed back on standard food and returned to the incubator at 29°C. Extravasation of the blue dye into the hemolymph was used as a reporter for increased IP as previously described.[16, 17] Smurfed flies were counted 24 h after exposure.

## References

1. Perouansky, M., General anesthetics and long-term neurotoxicity. Handb Exp Pharmacol 2008, (182), 143–57.

2. Kayser, E. B.; Morgan, P. G.; Sedensky, M. M., GAS-1: A mitochondrial protein controls sensitivity to volatile anesthetics in the nematode Caenorhabditis elegans. Anesthesiology 1999, 90, (2), 545–554.

3. Kayser, E. B.; Morgan, P. G.; Sedensky, M. M., Mitochondrial complex I function affects halothane sensitivity in Caenorhabditis elegans. Anesthesiology 2004, 101, (2), 365–72.

4. Morgan, P. G.; Hoppel, C. L.; Sedensky, M. M., Mitochondrial defects and anesthetic sensitivity. Anesthesiology 2002, 96, (5), 1268–70.

5. Niezgoda, J.; Morgan, P. G., Anesthetic considerations in patients with mitochondrial defects. Paediatric anaesthesia 2013, 23, (9), 785–93.

6. Finsterer, J., Leigh and Leigh-like syndrome in children and adults. Pediatr Neurol 2008, 39, (4), 223–35.

7. Quintana, A.; Kruse, S. E.; Kapur, R. P.; Sanz, E.; Palmiter, R. D., Complex I deficiency due to loss of Ndufs4 in the brain results in progressive encephalopathy resembling Leigh syndrome. Proceedings of the National Academy of Sciences of the United States of America 2010, 107, (24), 10996–1001.

8. Loewen, C. A.; Ganetzky, B., Mito-Nuclear Interactions Affecting Lifespan and Neurodegeneration in a Drosophila Model of Leigh Syndrome. Genetics 2018, 208, (4), 1535–1552.

9. Xu, H.; DeLuca, S. Z.; O’Farrell, P. H., Manipulating the metazoan mitochondrial genome with targeted restriction enzymes. Science 2008, 321, (5888), 575–7.

10. Olufs, Z. P. G.; Loewen, C. A.; Ganetzky, B.; Wassarman, D. A.; Perouansky, M., Genetic variability affects absolute and relative potencies and kinetics of the anesthetics isoflurane and sevoflurane in Drosophila melanogaster. Scientific reports 2018, 8, (1), 2348.

11. Olufs, Z. P. G.; Ganetzky, B.; Wassarman, D. A.; Perouansky, M., Mitochondrial Complex I Mutations Predispose Drosophila to Isoflurane Neurotoxicity. Anesthesiology 2020, 133, (4), 839–851.

12. Sazanov, L. A., A giant molecular proton pump: structure and mechanism of respiratory complex I. Nature reviews. Molecular cell biology 2015, 16, (6), 375–88.

13. Kirby, D. M.; Crawford, M.; Cleary, M. A.; Dahl, H. H.; Dennett, X.; Thorburn, D. R., Respiratory chain complex I deficiency: an underdiagnosed energy generation disorder. Neurology 1999, 52, (6), 1255–64.

14. Weaver, J. L., The brain-gut axis: A prime therapeutic target in traumatic brain injury. Brain Res 2021, 1753, 147225.

15. Katzenberger, R. J.; Chtarbanova, S.; Rimkus, S. A.; Fischer, J. A.; Kaur, G.; Seppala, J. M.; Swanson, L. C.; Zajac, J. E.; Ganetzky, B.; Wassarman, D. A., Death following traumatic brain injury in Drosophila is associated with intestinal barrier dysfunction. eLife 2015, 4.

16. Rera, M.; Bahadorani, S.; Cho, J.; Koehler, C. L.; Ulgherait, M.; Hur, J. H.; Ansari, W. S.; Lo, T., Jr.; Jones, D. L.; Walker, D. W., Modulation of longevity and tissue homeostasis by the Drosophila PGC-1 homolog. Cell metabolism 2011, 14, (5), 623–34.

17. Rera, M.; Clark, R. I.; Walker, D. W., Intestinal barrier dysfunction links metabolic and inflammatory markers of aging to death in Drosophila. Proceedings of the National Academy of Sciences of the United States of America 2012, 109, (52), 21528–33.

18. Njoku, D.; Laster, M. J.; Gong, D. H.; Eger, E. I., 2nd; Reed, G. F.; Martin, J. L., Biotransformation of halothane, enflurane, isoflurane, and desflurane to trifluoroacetylated liver proteins: association between protein acylation and hepatic injury. Anesth Analg 1997, 84, (1), 173–8.

19. Sawyer, D. C.; Eger, E. I., 2nd; Bahlman, S. H.; Cullen, B. F.; Impelman, D., Concentration dependence of hepatic halothane metabolism. Anesthesiology 1971, 34, (3), 230–5.

20. Miller, R. N.; Hunter, F. E., Jr., The effect of halothane on electron transport, oxidative phosphorylation, and swelling in rat liver mitochondria. Molecular pharmacology 1970, 6, (1), 67–77.

21. Ross, W. T., Jr.; Daggy, B. P.; Cardell, R. R., Jr., Hepatic necrosis caused by halothane and hypoxia in phenobarbital-treated rats. Anesthesiology 1979, 51, (4), 327–33.

22. Quintana, A.; Morgan, P. G.; Kruse, S. E.; Palmiter, R. D.; Sedensky, M. M., Altered anesthetic sensitivity of mice lacking Ndufs4, a subunit of mitochondrial complex I. PloS one 2012, 7, (8), e42904.

23. Falk, M. J.; Kayser, E. B.; Morgan, P. G.; Sedensky, M. M., Mitochondrial complex I function modulates volatile anesthetic sensitivity in C. elegans. Current biology : CB 2006, 16, (16), 1641–5.

24. Kirkman, M. A.; Yu-Wai-Man, P.; Korsten, A.; Leonhardt, M.; Dimitriadis, K.; De Coo, I. F.; Klopstock, T.; Chinnery, P. F., Gene-environment interactions in Leber hereditary optic neuropathy. Brain : a journal of neurology 2009, 132, (Pt 9), 2317–26.

25. Acharya, N. K.; Goldwaser, E. L.; Forsberg, M. M.; Godsey, G. A.; Johnson, C. A.; Sarkar, A.; DeMarshall, C.; Kosciuk, M. C.; Dash, J. M.; Hale, C. P.; Leonard, D. M.; Appelt, D. M.; Nagele, R. G., Sevoflurane and Isoflurane induce structural changes in brain vascular endothelial cells and increase blood-brain barrier permeability: Possible link to postoperative delirium and cognitive decline. Brain research 2015, 1620, 29–41.

26. Burman, J. L.; Itsara, L. S.; Kayser, E. B.; Suthammarak, W.; Wang, A. M.; Kaeberlein, M.; Sedensky, M. M.; Morgan, P. G.; Pallanck, L. J., A Drosophila model of mitochondrial disease caused by a complex I mutation that uncouples proton pumping from electron transfer. Disease models & mechanisms 2014, 7, (10), 1165–74.

